# Kinetic tracking of *Plasmodium falciparum* antigen presentation reveals determinants of protein export and membrane insertion

**DOI:** 10.1101/2021.09.06.459085

**Authors:** Jinfeng Shao, Gunjan Arora, Javier Manzella-Lapeira, Joseph A. Brzostowski, Sanjay A. Desai

**Author notes:** Address correspondence to Sanjay A. Desai.

## Abstract

Intracellular malaria parasites export many proteins into their host cell, inserting several into the erythrocyte plasma membrane to enable interactions with their external environment. While static techniques have identified some surface-exposed proteins, other candidates have eluded definitive localization and membrane topology determination. Moreover, both export kinetics and the mechanisms of membrane insertion remain largely unexplored. We introduce Reporter of Insertion and Surface Exposure (RISE), a method for continuous nondestructive tracking of antigen exposure on infected cells. RISE utilizes a small 11 aa NanoLuc fragment inserted into a target protein and detects surface exposure through high-affinity complementation. We tracked insertion of CLAG3, a malaria parasite protein linked to nutrient uptake, throughout the *P. falciparum* cycle in human erythrocytes. Our approach also revealed key determinants of trafficking and surface exposure. Removal of a C-terminal transmembrane domain aborted export. Unexpectedly, certain increases in the exposed reporter size improved surface exposure by up to 50-fold, revealing that both size and charge of the extracellular epitope influence membrane insertion. Insertion of parasite proteins at the host cell surface and antigen accessibility is regulated by multiple factors, enabling intracellular parasite survival and immune evasion under a broad range of conditions.

## Introduction

Many viral, bacterial and parasitic microbes invade, grow, and replicate within host cells to evade immune detection, access host cell machinery for replication, and use cellular macromolecules as nutrient sources [1]. At the same time, the intracellular milieu limits the pathogen from accessing plasma nutrients and often provides an inhospitable ionic composition or acidic pH. To overcome these hurdles, intracellular pathogens often export effector proteins into their host cell to remodel their abode, altering host cell defenses and physiology to their benefit [2–4]. A subset of effector proteins then insert in the host membrane to enable pathogen interactions with the extracellular space. These exposed proteins serve diverse roles and are acknowledged vaccine and drug targets.

In the virulent human malaria parasite *Plasmodium falciparum*, surface-exposed proteins benefit pathogen replication by facilitating cytoadherence [5], immune evasion [6] and nutrient uptake [7]. These antigens have been identified through static assays such as confocal and electron microscopy techniques. Because these methods lack the required spatial resolution [8], they cannot unambiguously determine if some parasite proteins are surface-exposed or only adherent to the inner membrane face or cytoskeleton. In some cases, susceptibility to extracellular proteases or antibody-based assays with live cells can resolve this uncertainty [9,10], but these approaches also have limitations. Another approach, mass spectrometry-based identification after surface labeling with NHS esters [11], has yielded a largely unvalidated list of proteins that is complicated by increased NHS ester permeability after infection with *Plasmodium* spp. [12].

Currently available methods also suffer from an inability to track the timing and kinetics of antigen insertion at the host membrane, hindering molecular insights. Both surface exposure via fusion of exocytic vesicles and “punch-through” insertion of soluble protein into the host membrane have been proposed [13,14], but the absence of direct and quantitative measurements has prevented definitive mechanistic insights.

To address these limitations and better define how pathogen proteins insert at their host cell membrane, we developed and used a Reporter of Insertion and Surface Exposure (RISE). Our study combines RISE with biochemical studies of the target reporter protein to identify constraints on protein trafficking and membrane insertion.

## Results

### HiBiT tagging within a conserved surface antigen in malaria parasites

We sought to generate a sensitive kinetic reporter for protein insertion on infected cells and chose human erythrocytes infected with the virulent *P. falciparum* malaria parasite. Indirect immunofluorescence microscopy assays (IFA) with antibodies against exposed epitopes offer a specific readout but suffer from low spatial resolution and provide limited kinetic information about protein export and host membrane insertion. Split enzyme reporters can overcome these limitations when one enzyme fragment is introduced into the exported protein; a specific signal is produced through complementation with a second fragment added extracellularly. Similar location-specific complementation has been described using mammalian proteins [15], but this approach has not been used to track appearance of pathogen-derived antigens on host cells. Because it relies on extracellular interactions with a surface-exposed epitope, our strategy resembles antigen presentation on immune effector cells [16].

The bright NanoLuc luciferase is an ideal enzymatic reporter for bloodstage *P. falciparum* studies [17] and has recently been optimized for development of a split reporter [18]. We selected the 11 residue HiBiT and 18 kDa LgBiT fragments of NanoLuc for our studies, based on their strong association (*K*_*D*_ = 700 pM) that yields an ATP-independent, furimazine-sensitive luminescence signal.

We next reasoned that minimal modification of a normally exported parasite protein would be more informative about parasite biology than extensively engineered reporters, as used previously [19]. We preferred the parasite CLAG3 protein for these studies over antigens such as PfEMP1 and RIFINs encoded by large multigene families to avoid epigenetic regulation and variable expression [20]. *Clag* paralogs also undergo epigenetic silencing [21], but some clones carry a single constitutively expressed hybrid *clag3* gene termed *clag3h* [22]. One such line, KC5, has been successfully used for transfections without the risk of epigenetic silencing [23]. CLAG3 is also the only known surface-exposed protein conserved in all examined *Plasmodium* spp. [24], suggesting that trafficking insights made using this protein may be broadly applicable. Because CLAG3 expression is linked to the plasmodial surface anion channel (PSAC), an ion and nutrient uptake channel at the host membrane [9], transport studies with transfectant parasites would also provide a biochemical correlate of reporter signal activity. We therefore selected CLAG3 for tagging and the KC5 clone for production of a surface exposure reporter parasite.

CLAG3 has a small 10-30 aa hypervariable region that appears to be exposed at the host membrane (HVR, Fig 1A) [25]. We therefore used CRISPR/Cas9 editing to replace the KC5 *clag3h* HVR sequence with a single HiBiT tag flanked by 8 aa linker sequences; we named this limiting dilution clone *8-1* based on the size of the flanking linker and the number of inserted HiBiT tags (Fig 1A, bottom). HVR replacement increased the size of the extracellular loop domain by a modest 10 residues (S1A Fig). We also produced *8-1HA*, a similar line with an HA epitope tag added after the HiBiT linker cassette. Two additional lines, *8-1trunc* and *8-1HAtrunc*, express truncated CLAG3 reporters with a stop codon introduced after the inserted cassette (Fig 1A, bottom and S1A Fig). We initially reasoned that these truncation constructs would yield a more flexible extracellular HiBiT epitope and provide insights into roles served by the downstream CLAG3 sequence. Although CLAG3 is a critical determinant of PSAC activity, CLAG3 knockout parasites are viable [26], possibly because other CLAG paralogs in *P. falciparum* compensate for CLAG3 loss. Thus, we expected that our modifications would be tolerated unless they produce a dominant-negative effect on nutrient uptake [27].

**Fig. 1.**
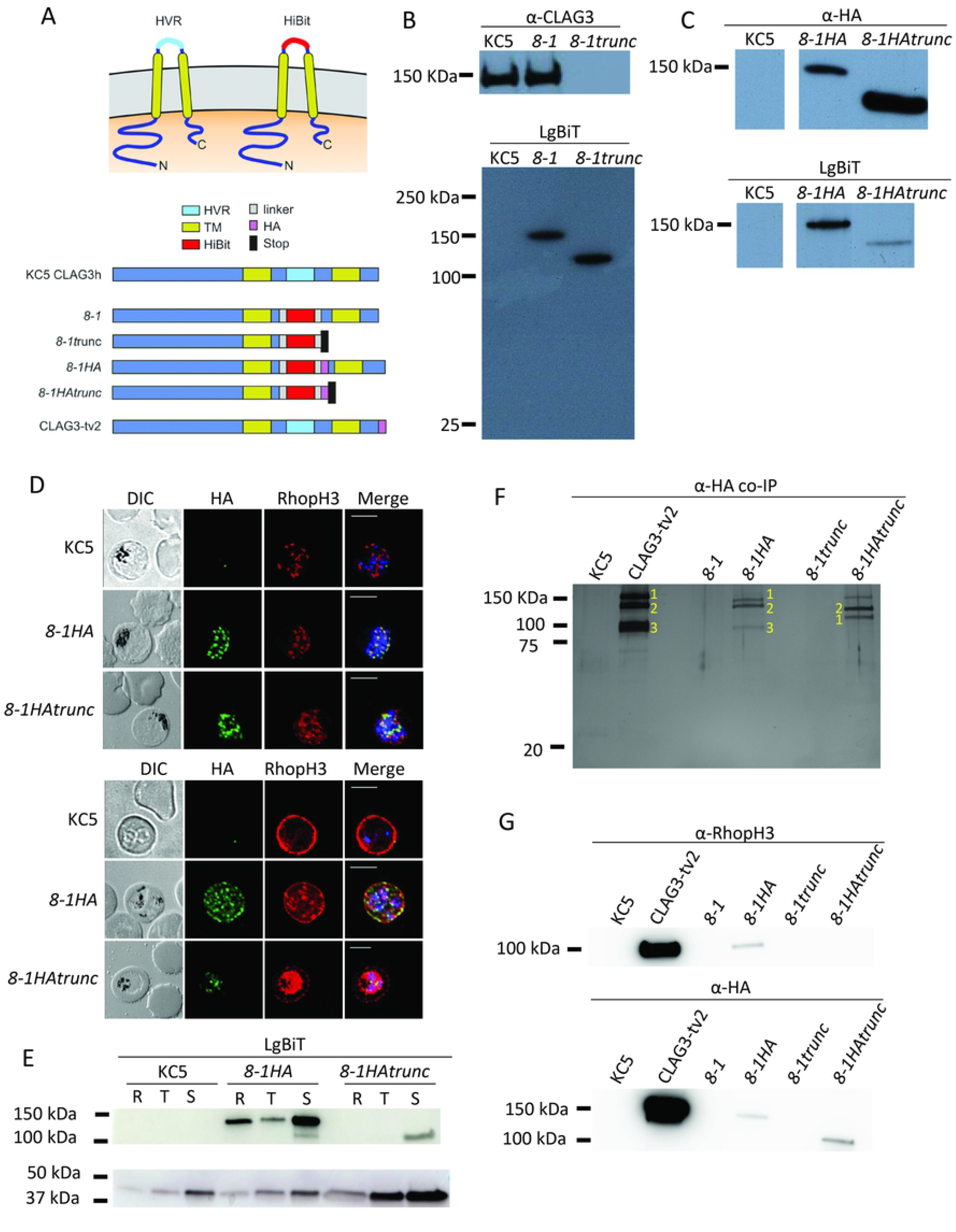
Design and production of host membrane-exposed split-reporter antigens. **(A)** Schematic shows native and modified CLAG3 topology at the host erythrocyte surface (left and right top images, respectively). The native HVR sequence is replaced by varied HiBiT reporter cassettes. Engineered lines are shown with ribbon diagrams at the bottom. (**B)** Immunoblots of matched total cell lysates from indicated lines, probed with antibody against a C-terminal CLAG3 epitope (top) or with LgBiT (bottom). *8-1trunc* is recognized by LgBiT but not by anti-CLAG3. **(C)** Immunoblots showing recognition of HA-tagged lines with anti-HA and LgBiT. **(D)** Indirect immunofluorescence assays (IFA) of indicated proteins in wild-type (WT) and transfected lines. In schizonts (top panels), CLAG3 colocalizes with RhopH3 in apical rhoptries (puncta) in *8-1HA*, but has a more diffuse distribution in *8-1HAtrunc*. IFA with trophozoite-stage parasites (bottom panels) reveals normal export of CLAG3 and colocalization with RhopH3 at the host membrane in *8-1HA* but a reduced signal with failed export in *8-1HAtrunc*. Scale bars, 5 mm. **(E)** Immunoblots showing stage-specific CLAG3 abundance in indicated lines (top row, probed with LgBiT). The truncated protein in *8-1HAtrunc* is detected in schizonts (S) but not rings or trophozoites (R and T). Bottom row, aldolase loading control. **(F)** Silver-stained gel showing co-immunoprecipitation using anti-HA beads and indicated parasite lysates. WT, *8-1*, and *8-1trunc* represent no-HA negative controls. Yellow 1, 2, and 3 labels indicate CLAG3, RhopH2, and RhopH3, respectively. **(G)** Immunoblots using eluates from panel **F**, probed with anti-RhopH3 and anti-HA for the CLAG3 bait protein.

Immunoblotting with each cloned transfectant confirmed expression and revealed single bands of expected size. Probing with anti-CLAG3 confirmed loss of this antibody’s C-terminal epitope in the *8-1trunc* clone and unchanged electrophoretic migration in *8-1* (Fig 1B, top). Both engineered CLAG3 isoforms were identified using a LgBiT probe that binds to HiBiT-tagged proteins to produce a luminescence signal (Fig 1B, bottom); the KC5 parent was not recognized by LgBiT, confirming specificity of this probe for the HiBiT tag. Similar results were obtained in the HA tandem-tagged parasites (Fig 1C), establishing faithful expression.

IFA confirmed and extended these findings. At the schizont stage, we detected each variant shortly after stage-specific synthesis under the genomic *clag3h* promoter (Fig 1D, upper group of images). While *8-1 HA* parasites trafficked the modified CLAG3 protein normally to developing rhoptries, the truncated tagged protein in *8-1HAtrunc* produced a more diffuse pattern with a small fraction reaching the rhoptry to colocalize with RhopH3, an associated protein that also contributes to PSAC formation [28].

At merozoite egress and reinvasion, rhoptry proteins are secreted into the next erythrocyte and deposited into the parasitophorous vacuole [29]. From there, through an incompletely understood interaction with the PTEX translocon, CLAG3 is exported into host cytosol for trafficking to the host membrane [28,30]. Imaging revealed that the tandem-tagged CLAG3 protein in *8-1HA* trafficked as expected and colocalized with RhopH3 at the host cell surface (Fig 1D, lower group); an antibody specific for the CLAG3 c-terminus further confirmed this localization (S1B Fig). In contrast, the truncated CLAG3 in *8-1HAtrunc* parasites was less abundant, suggesting that its poor trafficking to rhoptries compromised delivery to the next erythrocyte upon reinvasion. The small pool of this protein delivered into trophozoites failed to be exported and did not colocalize with RhopH3 (Fig 1D, bottom row).

Stage-specific immunoblotting using *8-1HA* parasites revealed increases in CLAG3 abundance upon parasite maturation to the schizont stage (Fig 1E), consistent with synthesis of this and other RhopH proteins predominantly in the late-stage parasites [31]. Ring and trophozoite parasites contained lower amounts that reflect incomplete transfer from prior cycle schizonts during egress and reinvasion [28]. The truncated protein in *8-1HAtrunc* was detected in schizonts but not in ring- and trophozoite-stage parasites, further implicating a role of the CLAG3 c-terminal region in efficient transfer to rhoptries and new erythrocytes during invasion.

We next used co-immunoprecipitation on anti-HA beads to examine protein-protein interactions for these CLAG3 reporter proteins (Fig 1F, silver-stained gel). CLAG3 was recovered from *8-1HA* and *8-1HAtrunc* lysates but not from negative control *8-1* and *8-1trunc* lines (bands labeled “1”), confirming specific pull-down. RhopH2 and RhopH3, unrelated proteins that interact with CLAG3 [24], were recovered from *8-1HA* (“2” and “3”), albeit with lower efficiency than in experiments using CLAG3-tv2, an engineered control parasite that a full-length CLAG3 with a C-terminal HA epitope tag (Fig 1A, ref # [30]). This reduced yield may result from compromised binding and recovery with an internal HA epitope tag when compared to the C-terminal tag in CLAG3-tv2. Co-immunoprecipitation using *8-1Hatrunc* yielded an unchanged RhopH2 band and a smaller band as expected for truncated CLAG3, but RhopH3 was not detected in these silver-stained gels (Fig 1F). Immunoblotting confirmed recovery of RhopH3 in *8-1HA* pull-downs and failed interaction with RhopH3 upon CLAG3 truncation (Fig 1G). The recent cryo-EM RhopH complex structure reveals that CLAG3 interacts with RhopH3 via two primary domains termed the CLAG3 “300 region” and “1300 loop” [30]. Because the 1300 loop is distal to the site of CLAG3 truncation in *8-1HAtrunc*, these findings suggest that this loop is required for stable CLAG3-RhopH3 interaction.

### Kinetics of membrane insertion and surface exposure

We next monitored stage-specific CLAG3 surface exposure on infected erythrocytes with the RISE method. We measured luminescence resulting from complementation of the HiBiT tag by extracellular LgBiT (Fig 2A). Bioluminescence microscopy revealed an undetectable reporter signal on immature ring-infected erythrocytes (Fig 2B, left panels) on KC5 and both HiBiT tagged lines, consistent with the appearance of PSAC activity on infected cells only after parasite maturation [32]. In contrast, trophozoite-infected cells exhibited a surface-distributed luciferase signal specific to *8-1* (Fig 2B-C). The *8-1trunc* parasite matured normally but failed to produce a surface signal.

**Fig. 2.**
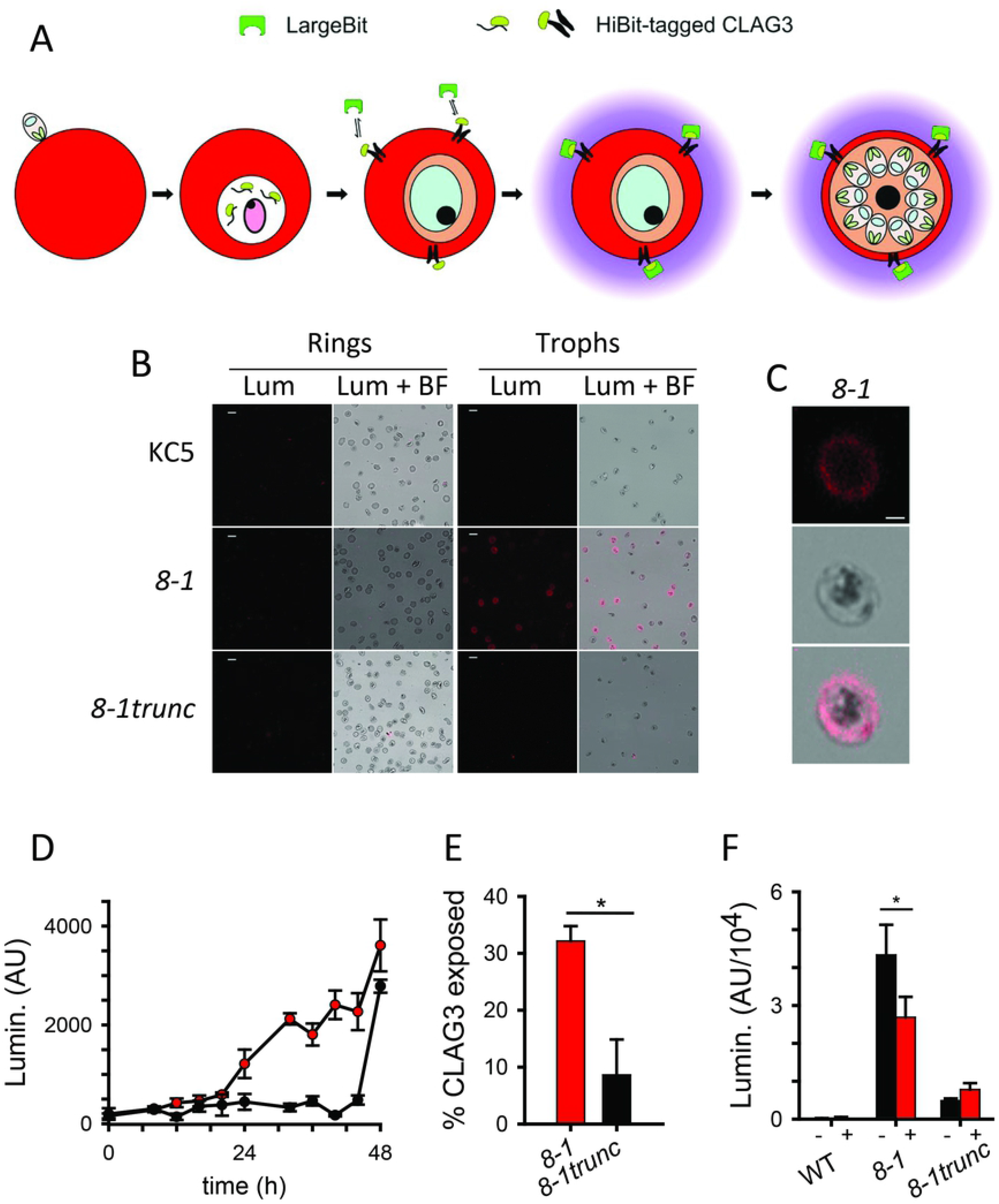
Faithful tracking of host membrane insertion. **(A)** Schematic showing parasite developmental stages and host membrane insertion-dependent luminescence. After invasion, the tagged CLAG3 protein is deposited into the parasitophorous vacuole before export and eventual insertion at the host membrane. Interaction between extracellular LgBiT and the surface-exposed HiBiT tag on *8-1* yields luminescence at mature parasite stages (purple glow). **(B)** Bioluminescence microscopy images showing undetectable signals on immature ring-infected cells (left panels), but a bright luminescence signal on *8-1* infected cells at the trophozoite stage (right panels). KC5 and *8-1trunc* parasites yield negligible signals. Scale bars, 10 µm. **(C)** “Zoom in” of a single *8-1* infected cell from panel **B**, showing surface distribution of luminescence signal. Scale bar, 2 µm. **(D)** Luminescence kinetics over parasite development, showing CLAG3 membrane insertion in *8-1* but not *8-1trunc* (red and black symbols, respectively; mean ± S.E.M. of 3 replicate wells, representative of 3 independent trials). Enriched synchronized early ring-infected cells seeded at *t* = 0. Increased signals at 48 h in both parasites reflect parasite egress and release of intracellular reporter protein. **(E)** Mean ± S.E.M. CLAG3 exposure on indicated lines at 36 h (*, *P* = 0.03, *n* = 3 trials), calculated as the luminescence signal normalized to total signal after cell lysis. **(F)** Mean ± S.E.M. luminescence signals from enriched trophozoite-infected cells without and with extracellular protease treatment (black and red bars, respectively; *, *P* = 0.005, *n* = 5).

We then miniaturized this reporter assay into 96-well microplate wells and tracked CLAG3 exposure kinetics in cultures initiated shortly after invasion. Although both clones exhibited negligible signals for the first 16 h of the parasite cycle, this lag was followed by a rapid increase in CLAG3 membrane insertion to produce luminescence on *8-1* (Fig 2D, red circles). This signal reached a plateau between 30 and 44 h on *8-1*, during which *8-1trunc* parasites continued to produce minimal luminescence. At the end of the erythrocyte cycle (44-48 h), both transfectant cultures exhibited abruptly increased signals, consistent with merozoite egress and CLAG3 discharge into extracellular medium [31]. At the signal plateau, approximately 1/3 of the CLAG3 within *8-1* infected cells had become surface-exposed based on measured reporter signal before and after detergent release (Fig 2E, 36 h timepoint).

Prior studies of the CLAG3 HVR implicated exposure at the host cell surface based on this motif’s susceptibility to extracellular protease [9,25]. We therefore examined whether the exposed HiBiT is also susceptible to external protease by measuring luminescence signals after a brief protease treatment of trophozoite infected cells. While the background signal in KC5 parasites and the low-level signal from *8-1trunc* cells were not significantly affected by extracellular protease treatment (*P* > 0.1, *n* = 5 independent trials each; Fig 2F), the large signal produced by *8-1* infected cells was reduced by 39 ± 3% upon treatment with extracellular protease (*P* = 0.005, *n* = 5), further confirming that our split NanoLuc assay faithfully reports on CLAG3 exposure at the host cell surface.

### Failed export compromises channel-mediated permeability

We next examined the effects of these CLAG3 modifications on nutrient uptake at the host membrane. We tracked uptake of sorbitol, a sugar alcohol with high PSAC permeability, and found that both the *8-1* and *8-1trunc* parasites increase host cell permeability (Fig 3A), as expected from its requirement for intracellular pathogen survival [27,28]. Both *8-1* and *8-1trunc* parasites exhibited lower sorbitol permeabilities than the parental KC5 (Fig 3B, *P* < 10_-4_, *n* = 20-21 trials each, one-way ANOVA with post-hoc tests), but uptake was preserved to a greater extent in *8-1*. The reduced permeability in *8-1trunc* matched that of a recently reported CLAG3 knockout, *C3h-KO* (*P =* 0.55; ref # [26]), indicating that the truncated CLAG3 in this parasite does not measurably contribute to PSAC activity. Failure to traffic and insert this protein in the host membrane conservatively accounts for this phenotype (Fig 2).

**Fig. 3.**
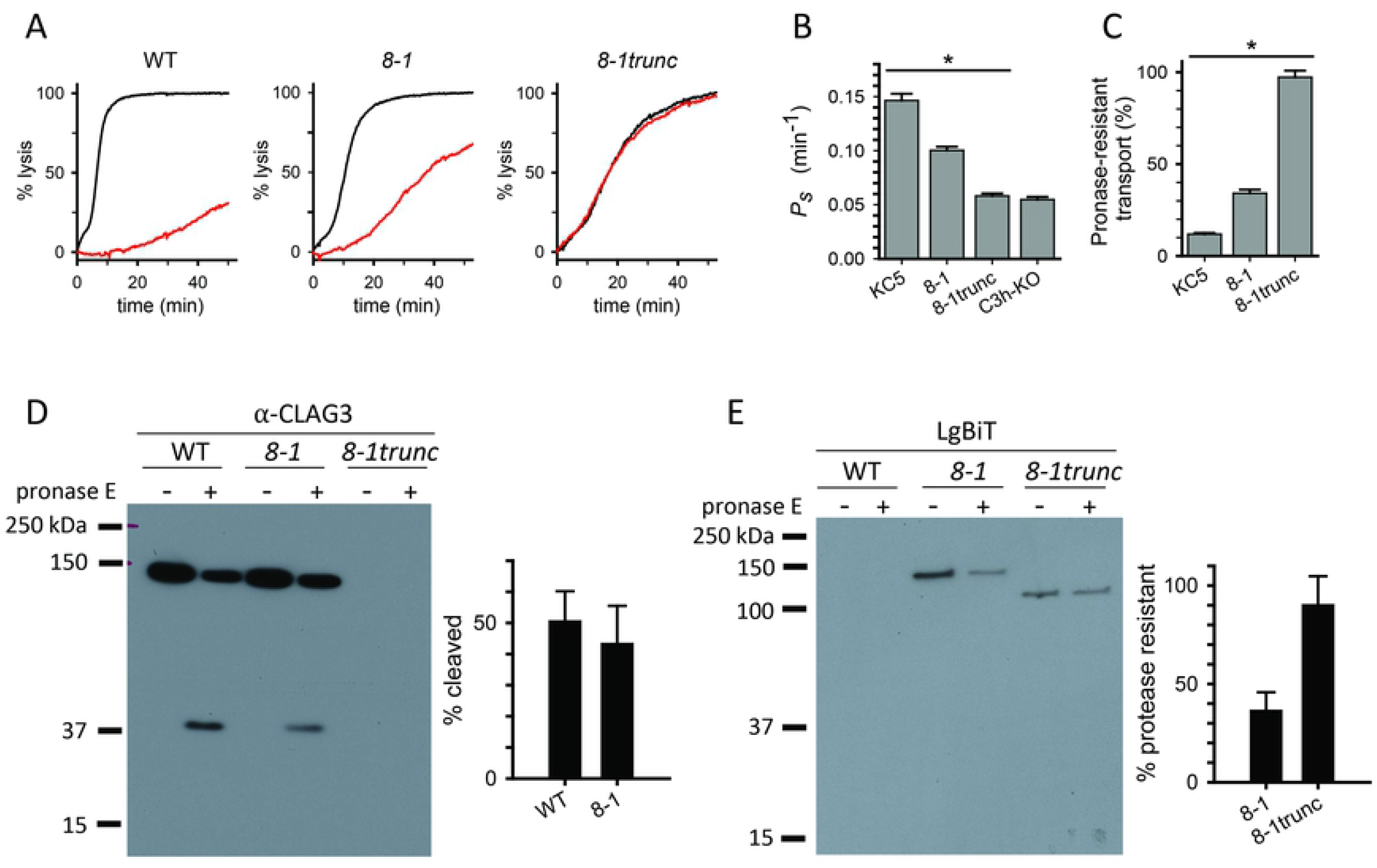
Modest effect of the reporter tag on CLAG3 export and on channel-mediated nutrient uptake. **(A)** Kinetics of osmotic lysis due to sorbitol uptake by indicated lines without or with pretreatment with pronase E (black and red traces, respectively). **(B)** Apparent sorbitol permeability coefficients from experiments as in panel A without protease treatment, calculated as 1/halftime of osmotic lysis. *, *P* < 10_-4_. **(C)** Pronase E-resistant transport activity for indicated lines, normalized to 100% for matched untreated controls. *, *P* < 10_-4_. **(D)** Anti-CLAG3 immunoblot showing a 37 kDa C-terminal cleavage product released by pronase E treatment. Bar graph shows quantified fractional band intensity of the cleavage product (mean ± S.E.M. 2 independent trials). **(E)** Blot probed with LgBiT showing reduction in the 150 kDa full length CLAG3 protein upon protease treatment. Bar graph, band intensities after protease treatment, normalized to 100% without protease (2 independent trials).

CLAG3 cleavage within the HVR by extracellular protease compromises solute transport [25]. Here, we found that pronase E treatment reduced channel-mediated transport in KC5 and *8-1*, but had no effect in *8-1trunc* parasites (red traces, Fig 3A; Fig 3C, *P* < 10_-4_, *n* = 10-11 trials, one way ANOVA with post-hoc tests), also consistent with intracellular retention of the truncated CLAG3 protein.

Immunoblots using an antibody directed against the CLAG3 C-terminus revealed single ∼ 37 kDa cleavage products in KC5 and *8-1* (Fig 3D), corresponding to proteolysis at the surface-exposed HVR and release of the distal fragment. Cleaved band intensities revealed that these two parasites exported and inserted CLAG3 protein at the host membrane with indistinguishable efficacies (Fig 3D, bar graph). Because *8-1trunc* parasites express a truncated CLAG3 not recognized by this antibody, we then probed these blots with LgBiT. This approach is based on visualization of HiBiT-tagged proteins via the luminescence generated upon LgBiT complementation. This treatment reduced the band intensity in protein from *8-1* parasites but had negligible effect on *8-1trunc* CLAG3 (Fig 3E, bar graph), consistent with proteolytic degradation of an exposed HiBit tag only on *8-1* parasites.

These studies establish that CLAG3 must insert at the host membrane to contribute to PSAC activity because the truncated protein in *8-1trunc* parasites has transport activity matching that of a CLAG3-null parasite. They also reveal that proteolysis at a surface-exposed loop on CLAG3 compromises transport regardless of sequence as this site retained its susceptibility when replaced by a HiBiT reporter.

### Membrane insertion not compromised by increased extracellular loop size

To examine possible constraints on insertion of the CLAG3 HVR at the host membrane, we generated an additional transfectant carrying a larger 3xHA epitope tag after the HiBiT cassette (S1C Fig). Biochemical studies with *8-1HA* and this new parasite, *8-1-3HA*, revealed marked increases in luminescence signals as the extracellular loop size increased by 9 and 27 residues, respectively. Using matched numbers of trophozoite-stage parasites, we found a 7- and 50-fold higher luminescence signals from the *8-1HA* and *8-1-3HA* parasites (Fig 4A, red bars) than from *8-1*. The increases in luminescence were more modest when measured after cell lysis with detergent (black bars). Greater accentuation with intact cells than after lysis is consistent with steric hindrance or constrained HiBiT presentation in a minimal extracellular loop, as proposed for CLAG3 [25]. Addition of these tags also significantly increased susceptibility of the HiBiT reporter to extracellular protease, with the larger 3xHA tag producing a greater reduction in luminescence upon protease treatment (Fig 4B). Membrane insertion and surface exposure were further confirmed with immunoblotting (Fig 4C) and luminescence imaging, which revealed dramatically increased signals from intact cells (Fig 4D). These findings suggest improved LgBiT binding and reporter complementation upon adjacent HA epitope tagging, presumably because the size and negative charge of this tag improves accessibility at the extracellular loop and within soluble RhopH complexes upon detergent release.

**Fig. 4.**
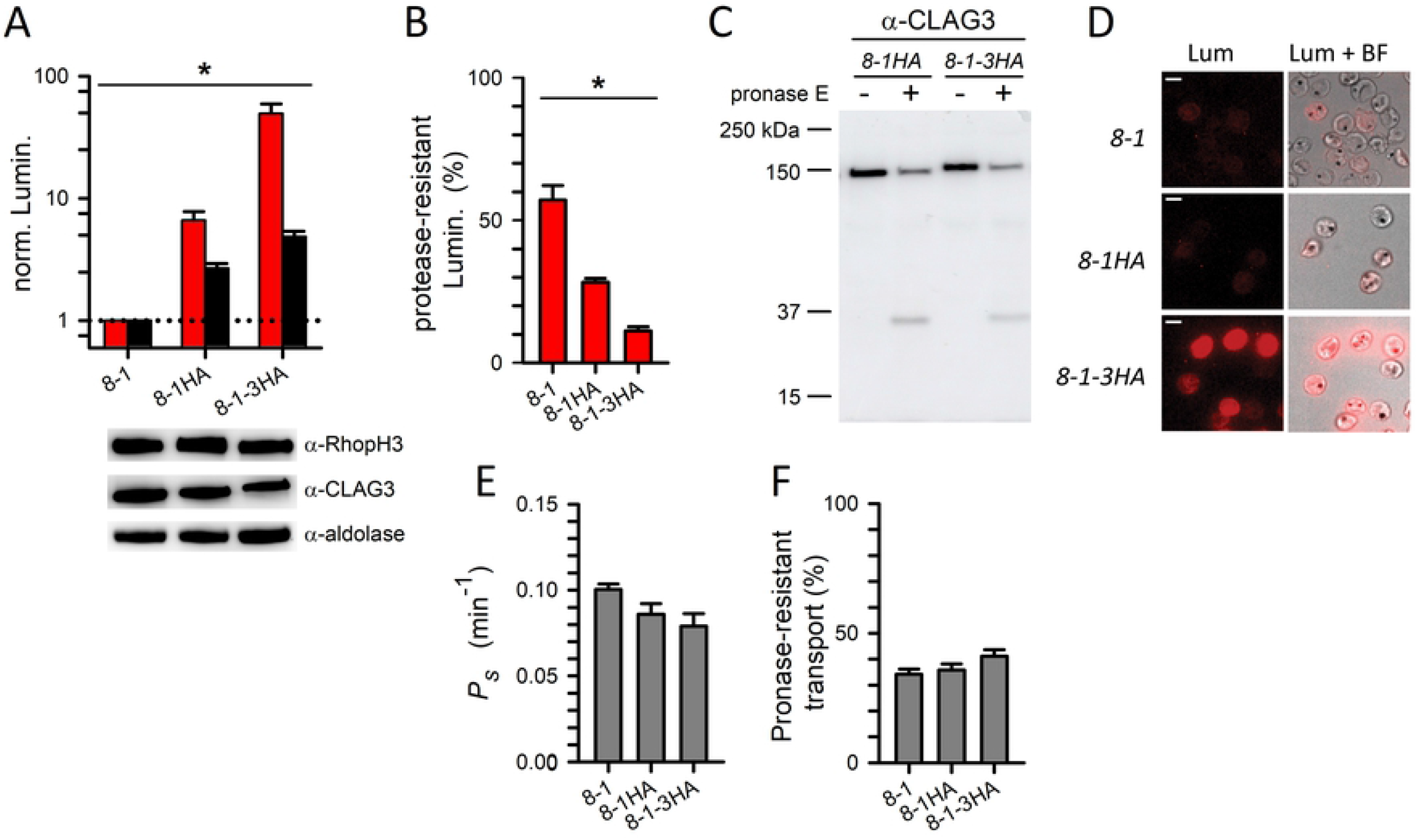
Addition of HA and 3xHA tags improve reporter signal and does not compromise host membrane insertion. **(A)** Mean ± S.E.M. luminescence signals from indicated parasite clones using matched amounts of enriched trophozoite-infected cells. Red and black bars represent intact and lysed cells, respectively (*, *P* = 10_-4_, one-way ANOVA, *n* = 3 independent trials). Signals are shown after normalization of *8-1* readings to 1.0 in each trial. Immunoblot shows representative loading control for matching protein contents from one of the three trials. **(B)** Mean ± S.E.M. luminescence signals remaining after extracellular protease treatment of intact cells from indicated lines, normalized to 100% for no protease effect (*, *P* = 0.0002, *n* = 3-6 for each clone). **(C)** Anti-CLAG3 immunoblot showing C-terminal cleavage product released by extracellular pronase E treatment. **(D)** Bioluminescence microscopy showing increased signals from individual cells when 1HA and 3HA tags are added to the HiBit reporter. Scale bars, 5 µm. **(E)** Mean ± S.E.M. sorbitol permeabilities for indicated parasites, determined from osmotic lysis experiments (*n* = 5-21 trials each). **(F)** % of transport resistant to treatment with extracellular pronase E (*n* = 4-11 trials). Channel-mediated transport is modestly affected in these lines.

These larger insertions into CLAG3 had modest effects on channel-mediated sorbitol uptake at the host membrane (Fig 4E). The resulting channels also retained quantitatively similar protease susceptibilities (Fig 4F), consistent with minimally affected CLAG3 insertion and PSAC formation at the host membrane.

### Larger epitopes reveal complex regulation of CLAG3 membrane insertion

To further explore size and charge constraints on pathogen epitope presentation, we made additional transfectants containing multiple HiBiT tags with two different linker sizes (Fig 5A). As these constructs retained upstream and distal CLAG3 sequences, each modified protein trafficked normally through schizonts and was delivered into maturing trophozoite-infected cells (S2A-B Fig), where the increased size of the targeted protein was apparent in immunoblots (Fig 5B). Although immunofluorescence, sorbitol permeability measurements and protease susceptibility studies all suggested that CLAG3 failed to export and undergo host membrane insertion to enable PSAC activity in these lines (S2B Fig and Fig 5C), bioluminescence intensity analyses using RISE identified individual cells that presented CLAG3 on host cells (Fig 5D). Notably, while most cells in each of the largest multiple HiBiT constructs produced background signals, a few cells produced very bright signals that exceeded those seen on *8-1* parasites. These intense signals may reflect either reduced steric hindrance with larger, more flexible extracellular loops or signal amplification from HiBiT multiplicity on each protein. The markedly differing signals from individual cells is unexpected for these clonal lines. This observation suggests epigenetic control of CLAG3 export and host membrane insertion.

**Fig. 5.**
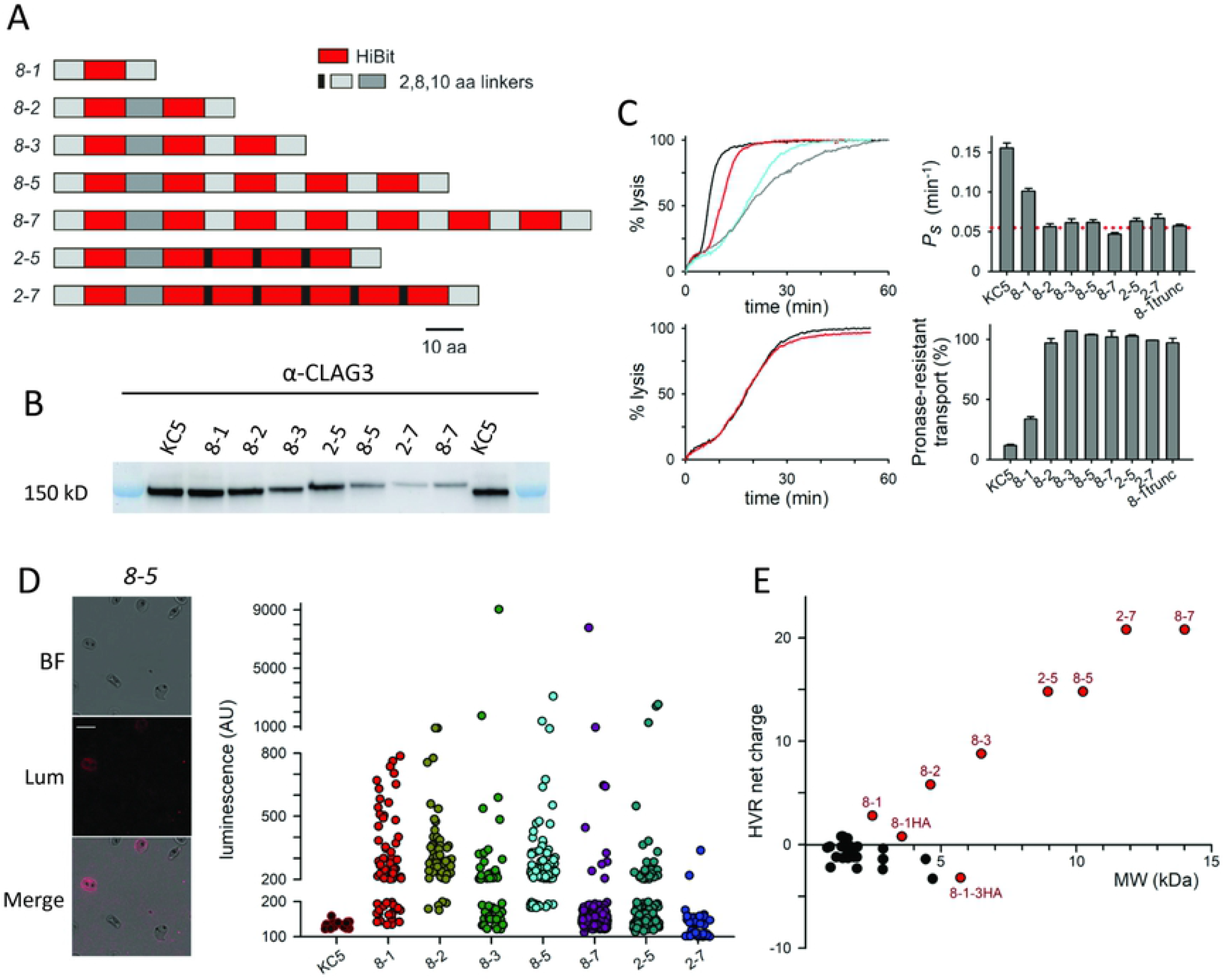
Large reporter inserts reveal complex regulation of CLAG3 insertion at the host membrane. **(A)** To-scale schematic showing multi-HiBiT inserts and sizes of linkers between elements. The parasite names reflect the predominant linker size (in residues) followed by the number of introduced HiBiTs. **(B)** Anti-CLAG3 immunoblot confirming the expected increase in CLAG3 size with each construct. **(C)** PSAC activity in transfected clones. Top left, osmotic lysis kinetics for the KC5 parent (black), *8-1* (red), *8-2* (blue), and *8-1trunc* (grey). Right, mean ± S.E.M. apparent sorbitol permeability coefficients for indicated lines; red dotted line, mean for the C*3h-KO* knockout line. Note that all multi-HiBiT inserts have permeabilities indistinguishable from those of the CLAG3 knockout. Bottom left, *8-2* osmotic lysis kinetics with and without protease treatment (red and black, respectively), showing that transport is not protease sensitive in this line. Bottom right, mean ± S.E.M. % of transport resistant to extracellular pronase E treatment. In contrast to the CLAG3 export competent KC5 and *8-1* lines, transport linked to the multi-HiBiT inserts is resistant to extracellular pronase. **(D)** Microscopy images showing rare luminescent cells in the *8-5* reporter line. Jitter plot shows single cell intensities of indicated lines. Break points were selected based on the range of intensities in the KC5 and *8-1* controls. Note that while most cells are negative in the multi-HiBiT constructs, a small number have very high signals in each reporter line. **(E)** 2D plot showing molecular weight (MW) and net charge at pH 7.4 for the CLAG3 HVR region from 38 available sequences (black circles) and the indicated engineered substitutions (red symbols).

We tabulated the HVR sequences from 38 available CLAG3 sequences and compared their properties to those of the lines we have engineered. Although they are variant, the native HVR sequences tended to have a modest net negative charge (Fig 5E, black symbols). In contrast, the constructs containing more than one HiBit epitope were increasingly basic, yielding net positive charges on the at the extracellular loop (red symbols). Notably, the sequence in *8-1-3HA*, whose CLAG3 successfully inserted at the host membrane to produce remarkably bright luminescence using our RISE assays, had a higher molecular weight than that of *8-2*, which failed to traffic CLAG3 protein faithfully in most cells. It appears that the domain’s net positive charge in *8-2* and other multiple HiBit lines prevents trafficking and host membrane insertion. Along with structural constraints that ensure CLAG3-mediated nutrient uptake and evolutionary pressures to evade host immunity, the extracellular loop of this conserved protein family must also meet charge and size requirements for faithful trafficking and host membrane insertion.

## Discussion

We present a new reporter that detects insertion of pathogen virulence antigens on their host cells. Our use of a split NanoLuc reporter is broadly applicable to a range of intracellular pathogens and will permit non-destructive kinetic tracking of antigens at the host cell surface. Some proteins targeted to underlying membranes, such as the parasitophorous vacuolar membrane of *Plasmodia, Toxoplasma*, and other parasites, may also be studied using selective permeabilization of the host membrane [33].

This reporter assay represents an important step toward understanding how pathogens interact with their host cells and will provide quantitative insights into the presentation of targeted antigens to the host immune system. We used this new technology to examine presentation of the conserved CLAG3 antigen on the surface of human erythrocytes. Associated nutrient channel transport, protein chemistry, and bioluminescence confocal microscopy studies all validated our new method.

Our findings implicate a revised model of CLAG3 trafficking. CLAG3 produced in schizonts remains inaccessible to extracellular LgBiT upon transfer to ring-infected cells; as the intracellular parasite matures, the protein is exported and inserts in the host membrane with kinetics that parallel the gradual appearance of the associated nutrient channel activity on trophozoite-infected cells. A C-terminal truncation that does not alter the reporter or its upstream sequence compromised trafficking, abolished host membrane insertion, and reduced host cell permeability to levels seen in a recently reported CLAG3-null parasite. Using varied reporter insertions, we also uncovered complexities in this protein’s insertion into the erythrocyte membrane; our studies suggest that both size and charge of the extracellular peptide loop determine host membrane insertion.

Our luminescence imaging studies revealed marked variation between cells in cloned multiple HiBit reporter lines, with some cells exhibiting high levels of CLAG3 exposure despite insertion of large reporter domains. This finding implicates an epigenetic, post-translational mechanism for regulating surface exposure. We propose that this may reflect altered expression of one or more parasite chaperone proteins in host cytosol [34] or post-translational modifications of CLAG3 [35,36]. Epigenetic control of antigen presentation on infected cells has, to our knowledge, not been proposed for any intracellular pathogens. This finding reveals the remarkable sophistication of malaria parasites in controlling their interactions with host plasma and further promotes immune evasion.

We envision that the quantitative and sensitive readout enabled by a small HiBiT epitope inserted at exposed antigen sites will unveil how pathogens modify their host cells while evading immune attack.

## Materials and Methods

### Parasite cultivation and transfection

The *P. falciparum* KC5 parasite clone and its engineered derivatives were cultivated in O+ human erythrocytes (Interstate Blood Bank, Inc.) at 5% hematocrit in standard RPMI 1640-based media (KD medical) supplemented with supplemented with 25 mM HEPES, 50 μg/mL hypoxanthine, 0.5% NZ Microbiological BSA (MP Biomedicals), gentamicin and 28.6 mM NaHCO_3_ (Gibco) at 37 _o_C under 90% N_2_, 5% CO_2_, 5% O_2_.

CRISPR-Cas9 DNA transfection of parasites to produce reporter lines was performed using electroporation of pUF1-Cas9 and modified pL6 plasmids into uninfected erythrocytes as described previously [28]. Plasmids were constructed using synthetic double-stranded DNA (Integrated DNA Technologies) and In-Fusion cloning (Takara) into the pL6 plasmid. Single guide RNAs (sgRNA), selected using on-, off- and paralog specificity scores [37], were also introduced using In-Fusion. After erythrocyte electroporation and addition of schizont-staged parasites, the culture was selected with 1.5 µM DSM1 and 2.5 nM WR99210. After parasite outgrowth and PCR confirmation of integration, limiting dilution cloning was performed for all transfectant lines. All experiments were performed with sequence-verified clones.

### Immunoblotting

Synchronous parasite cultures were harvested, percoll-enriched where indicated, and used for immunoblotting experiments after hypotonic lysis (7.5 mM Na_2_HPO_4_, 1 mM EDTA, 1 mM PMSF, pH 7.5) and solubilization in Laemmli sample buffer containing 6% SDS. When required, samples were matched with turbidity measurements at 700 nm. Proteins were separated by SDS-PAGE (4–15% Mini-PROTEAN TGX gel, Bio-RAD) and transferred to nitrocellulose membranes. After blocking with 3% skim milk powder in 150 mM NaCl, 20 mM TrisHCl, pH 7.4 with 0.1% Tween20 at RT for 1 h, primary antibodies were applied in the same blocking buffer at a 1:1000-1:3000 dilution and incubated overnight at 4 °C with gentle rocking. After three washes in 150 mM NaCl, 20 mM TrisHCl, pH 7.4 with 0.1% Tween20, HRP-conjugated secondary antibodies were added at 1:3000 dilution. The blot was incubated for 1 h and washed three times. Imaging was performed after addition of Clarity Western ECL substrate (Bio-Rad) using the AI 680 imager (GE healthcare) or standard x-ray film exposure.

Where used, protease treatment was performed after washing and resuspending enriched trophozoite-infected cells in PBS-2 (150 mM NaCl, 20 mM Na_2_HPO_4_, 0.6 mM CaCl_2_, 1 mM MgCl_2_, pH 7.4) at 5% hematocrit with 1 mg/mL pronase E (Sigma) for 45-60 min at 37 _o_C. The treated cells and matched untreated cells were washed in PBS-2 with 1 mM PMSF at 4 _o_C before an additional wash in this buffer with 1 mM EDTA. After hypotonic lysis, the membrane fraction was harvested by ultracentrifugation (100,000 ×*g*, 1 h at 4 °C) and solubilized in Laemmli sample buffer as above.

Blots probed with LgBiT used the HiBiT blotting system kit (Promega). After protein transfer, nitrocellulose membranes were washed in 150 mM NaCl, 20 mM TrisHCl, pH 7.4 with 0.1% Tween20. LgBiT was applied in this blotting buffer at 1:200 dilution and incubated overnight at 4 °C with gentle rocking. Furimazine was then added in blotting buffer at a 1:500 dilution before imaging as above.

Band intensities were quantified using Image J software. Statistical analyses were based on three independent trials.

### Coimmunoprecipitation

Schizont-stage infected cells were percoll-sorbitol enriched and lysed with 20 volumes of 10 mM Tris pH 7.5, 300 mM NaCl, 1% Triton-X100, 1 mM PMSF. After a 30 min incubation at 4 °C, solubilized proteins were separated by centrifugation (14,000 x *g*, 15 min, 4°C) and incubated with anti-HA affinity agarose beads (Sigma) with gentle mixing overnight at 4°C. After five washes, bound protein was eluted by addition of 2.5 mg/mL HA peptide in 10 mM Tris pH 7.5, 250 mM NaCl, 0.1% Triton-X100 for 30 min. Eluted proteins were resuspended in Laemmli sample buffer and subjected to SDS-PAGE.

### Immunofluorescence assays

Indirect immunofluorescence assays were performed using air-dried thin smears after fixation with 1:1 acetone:methanol at -20 °C for 2 min. Slides were then dried, blocked with 3% milk in PBS for 1 h at RT, and incubated with primary antibodies in blocking buffer (mouse anti-CLAG3, 1:100; rabbit anti-RhopH3, 1:500; mouse anti-HA, 1:100) for 1.5 h at RT under coverslips. After two washes with chilled PBS, Alexa Fluor 488 or 594-conjugated secondary antibody at a 1:500 dilution and 10 µg/mL DAPI were added in blocking buffer and incubated for 30 min at RT. After washes and drying, slides were mounted with Prolong Diamond anti-fade mountant (Molecular Probes). Images were collected on a Leica SP8 microscope using a 64x oil immersion objective with serial 405 nm, 488 nm, or 594 nm excitation. Images were processed using Leica LAS X and Huygens software.

### Enrichment of ring-infected cells

Ring-stage cultures were further synchronized by a 20 min incubation in 4% xylitol, a sugar alcohol with high PSAC permeability [38], to lyse mature infected cells. The culture was then resuspended and incubated in culture medium supplemented with 207 mM xylitol for 1 h at 37 _o_C. This cell suspension was then layered on a discontinuous percoll-xylitol gradient a bottom layer of 72% Percoll and an upper layer of 40% Percoll; both solutions were prepared in RPMI 1640 medium with 208 mM xylitol, 12.4 mM HEPES, and 16.3 mg/L BSA. After centrifugation (10,000 x *g* for 30 min at 21 _o_C), ring-stage infected cells at 65-90% parasitemia were harvested and washed by dropwise addition of culture medium.

### Luminescence measurements and export kinetics

Luminescence measurements were performed using percoll-enriched infected erythrocytes in 384 well microplates using the Nano-Glo HiBiT extracellular detection system (Promega). Cells were resuspended at 0.2% hematocrit in culture medium diluted with two volumes of 200:1:50 buffer:LgBiT:Furimazine, according to the manufacture’s protocol. After a 30 min RT incubation, luminescence was measured using the Centro XS3 LB 960 reader (Berthold) or Synergy Neo2 (BioTek) with a counting time of 0.5 s/well.

CLAG3 export kinetics were tracked using luminescence after seeding enriched ring-infected cells in culture medium at 0.5% hematocrit into triplicate wells at 60 µL/well. The plates were sealed with Breathe-Easy sealing membrane (RPI) and incubated at 37 _o_C under 5% CO_2_ in air. At timed intervals, 40 µL of medium from selected wells was replaced with Nano-Glo HiBiT extracellular buffer with LgBiT and furimazine substrate (Promega). Readings were taken after a 30 min room temperature incubation as described above.

### Bioluminescence microscopy

Erythrocytes infected with reporter parasite clones were imaged using a LV200 inverted bioluminescence microscope with a temperature-controlled stage (Olympus). Enriched ring- or trophozoite-infected erythrocytes were resuspended in culture medium at 2.5% hematocrit before diluting 50x into Nano-Glo HiBiT extracellular buffer (Promega) with LgBiT and furimazine at 100x and 50x dilutions, respectively for a total volume of 100 µL in a 35 mm poly-D-lysine coated coverslip dish (MatTek). Cells were allowed to settle at 37 _o_C for 25 min in the LV200 microscope before selecting fields of view for imaging. Bioluminescence images were collected with a 45 min exposure under a 64x oil immersion objective. Images were visualized and adjusted to 14 bit in LCmicro_2.2 software (Olympus).

Single cell luminescence intensities were quantified using a locally-developed macro that uses the corresponding brightfield image to define the cell boundary. This macro reports luminescence intensities over the cell, tabulating mean, max and min values along with the cell 2D area and is available upon request.

### Osmotic lysis assays

The kinetics of PSAC-mediated sorbitol uptake and infected cell osmotic lysis were continuously tracked as described previously [26]. Enriched trophozoite-stage infected cells were washed and resuspended in 150 mM NaCl, 20 mM Na-HEPES buffer, 0.1 mg/mL BSA, pH 7.4. Solute uptake was initiated by the addition of 280 mM sorbitol, 20 mM Na-HEPES, 0.1 mg/mL BSA, pH 7.4. Concomitant uptake of sorbitol and water produces osmotic lysis at rates directly proportional to PSAC sorbitol permeability. Lysis kinetics were continuously monitored using transmittance of 700 nm light through the cell suspension. Osmotic lysis half-times, normalized permeability estimates, and measures of protease effect were determined from the recordings with locally developed code.

### Computational and statistical analyses

CLAG3 sequences were downloaded from www.plasmodb.org and aligned using Multiple Sequence Alignment (MUSCLE) to identify HVR sequences from available *P. falciparum* paralogs. The molecular weight and net charge at pH 7.4 was calculated at http://protcalc.sourceforge.net/.

Numerical data are shown as mean ± S.E.M. Data were analyzed in SigmaPlot 10.0 (Systat) or Prizm 8 (GraphPad). Statistical significance was determined using unpaired or paired Student’s *t-*test or one-way ANOVA with *post hoc* Tukey’s multiple comparisons test as appropriate. Significance was accepted at *P* < 0.05%.

## Acknowledgements

We thank Gagan Saggu for help with immunofluorescence assays and David Jacobus for WR99210. DSM1 (MRA-1161) was obtained through MR4 as part of the BEI Resources Repository, NIAID, NIH.

## Author contributions

### Conceptualization

Jinfeng Shao, Sanjay A. Desai.

### Data curation

Jinfeng Shao, Gunjan Arora, Javier Manzella-Lapeira, Joseph A. Brzostowski, Sanjay A. Desai.

### Formal analysis

Jinfeng Shao, Gunjan Arora, Javier Manzella-Lapeira, Joseph A. Brzostowski, Sanjay A. Desai.

### Investigation

Jinfeng Shao, Gunjan Arora, Javier Manzella-Lapeira.

### Supervision

Sanjay A. Desai.

### Writing – original draft

Jinfeng Shao, Sanjay A. Desai.

### Writing – review & editing

Jinfeng Shao, Gunjan Arora, Javier Manzella-Lapeira, Joseph A. Brzostowski, Sanjay A. Desai.

## Supporting Figure Legends

**S1 Fig. Construction and validation of reporter protein for host membrane insertion. (A)** Sequences of the modified CLAG3 locus in *8-1* and *8-1HA* reporter lines aligned with the parental KC5 CLAG3h and CLAG3.1 and CLAG3.2 from divergent *P. falciparum* lines (7G8 and Dd2 from Brazil and Indochina, respectively). The HVR sequences from wild-type lines is highlighted in grey; introduced HiBiT is shown with magenta highlight. HA tag is underlined; the sites where stop codons were introduced to produce *8-1trunc* and *8-1HAtrunc* are marked with red arrows. Identical and conserved residues that flank the HVR are in red and blue, respectively. Note the length polymorphism in native HVR sequences; the modification to produce *8-1HA* increases the length of this extracellular motif. **(B)** IFA of trophozoite-stage parasites probed with a CLAG3-specific antibody directed against a C-terminal epitope [9]. Scale bar, 5 µm. Notice unchanged export and colocalization with RhopH3 in *8-1* and *8-1HA* parasites. **(C)** Modified CLAG3 locus sequence in the *8-1-3HA* parasite with color-coding as in panel **A**. The 3xHA tag is underlined.

**S2 Fig. Trafficking and membrane insertion for large multi-HiBiT insertions into the CLAG3 HVR.** IFA micrographs of indicated lines at the schizont and trophozoite stages (panels **A** and **B**, respectively), showing that each construct is faithfully trafficked to rhoptry organelles in schizonts, but that the multi-HiBiT lines do not export CLAG3 to the host membrane of most trophozoite-infected cells. Scale bars, 5 µm. **(C)** Anti-CLAG3 immunoblot showing that chymotrypsin releases a ∼ 35 kDa cleavage product in KC5 and *8-1* parasites, but that cleavage is not detected in studies with the indicated multi-HiBiT lines.

